# Temporal proteomic profiling of postnatal human cortical development

**DOI:** 10.1101/188565

**Authors:** Michael S. Breen, Sureyya Ozcan, Jordan M. Ramsey, Zichen Wang, Avi Ma’ayan, Nitin Rustogi, Michael G. Gottschalk, Maree J. Webster, Cynthia Shannon Weickert, Joseph D. Buxbaum, Sabine Bahn

## Abstract

Healthy cortical development depends on precise regulation of transcription and translation. However, the dynamics of how proteins are expressed, function and interact across postnatal human cortical development remain poorly understood. We surveyed the proteomic landscape of 69 dorsolateral prefrontal cortex samples across seven stages of postnatal life and integrated these data with paired transcriptome data. We detected 911 proteins by liquid chromatography-mass spectrometry, and 83 were significantly associated with postnatal age (FDR *p* < 0.05). Network analysis identified three modules of co-regulated proteins correlated with age, including two modules with increasing expression involved in gliogenesis and NADH-metabolism and one neurogenesis-related module with decreasing expression throughout development. Integration with paired transcriptome data revealed that these age-related protein modules overlapped with RNA modules and displayed collinear developmental trajectories. Importantly, RNA expression profiles that are dynamically regulated throughout cortical development display tighter correlations with their respective translated protein expression compared to those RNA profiles that are not. Moreover, the correspondence between RNA and protein expression significantly decreases as a function of cortical aging, especially for genes involved in myelination and cytoskeleton organization. Finally, we used this data resource to elucidate the functional impact of genetic risk loci for intellectual disability, converging on gliogenesis, myelination and ATP-metabolism modules in the proteome and transcriptome. We share all data in an interactive, searchable companion website. Collectively, our findings reveal dynamic aspects of protein regulation and provide new insights into brain development, maturation and disease.

## Introduction

The human prefrontal cortex plays a critical role for higher cognitive processes, including executive function, social cognition and judgment, and has been implicated in the onset and progression of many, if not most, neurodevelopmental disorders^1-6^. The development of a properly functioning prefrontal cortex depends upon the proliferation and signaling of several cell types as well as the reprograming of transcriptional and translational pathways that unfold over the first two-three decades of postnatal life^7^. During this time, the brain quadruples in size, and grows through interneuronal genesis and maturation, glial multiplication, myelination, formation of new synaptic connections and pruning of unused synaptic connections^8-10^. Such processes are orchestrated by thousands of molecules in a tightly synchronized spatiotemporal fashion, and the disruption of any one of which may result in loss of cortical integrity and homeostasis^10^, leading to cognitive deficits seen in patients with neurodevelopmental abnormalities. Therefore, understanding the molecular factors governing long-term brain development in normal individuals is critical for the identification of neurodevelopment mechanisms and developmental vulnerability periods.

Much of our current knowledge of the biological changes underlying human brain development has been inferred from large transcriptomic investigations. Initial reports of the developing human brain transcriptome revealed marked changes across development and aging, with the largest gene expression changes occurring prenatally and during infancy and early childhood^11-14^; ages when many neurodevelopmental disorders become clinically recognizable. In parallel, several studies have identified developmental transcriptional networks with regional and cell type specific expression patterns enriched within neurodevelopmental disorder-associated genetic risk loci^15-18^, providing mechanistic insights of how mutations in risk genes might perturb typical brain development. Several studies consistently report however, that levels of messenger RNA and their respective translated proteins often correlate poorly^19-22^. As proteins are the main functional components in all cells, generating equivalent proteomic information across human brain development represents a critical gap in the field.

Mass spectrometry-based proteomics provides a comprehensive and complimentary perspective to transcriptomic changes and can serve as an indicator for functional and network levels of aging. Although human proteome research has predominately focused on defining a disease signature within a specific developmental period, there has been some progress in understanding the developmental proteome^23-25^. For example, a recent study profiled the orbitofrontal cortex and identified 127 proteins implicated in cellular growth and proliferation that were differentially expressed between young or old human male individuals^23^. A separate study profiled seven different brain regions across 11 developmentally distinct individuals and found substantial differences in protein abundance between brain regions, reflective of cytoarchitectural and functional variation^25^. While these investigations have been key for informing mechanisms of brain development, it has been challenging for proteome research to identify highly abundant and reproducible proteins across dozens of biological replicates. More selective mass spectrometry techniques that are tailored to detect highly abundant proteins across larger sample sizes are required to accurately infer long-term time-dependent protein expression patterns. As such, a critical remaining question is how the brain proteome unfolds throughout development, and ultimately how this information may inform brain mechanisms governing health and disease.

The current investigation applied label-free liquid chromatography-mass spectrometry (LC-MS) proteomics to 69 dorsolateral prefrontal cortex (DLPFC) samples from healthy individuals aged 39 days to 49.5 years. These proteomic data were integrated with paired transcriptome data from matching DLPFC samples and together comprise a unique resource of well-annotated anatomical structures of fresh human brains from seven different developmental stages. A multistep analytic approach was used that specifically sought to address two main goals: (1) to identify proteins and networks of highly correlated proteins significantly associated with distinct developmental stages and that changed with human age; and (2) to determine the biological organization of the proteome across postnatal development and clarify the relationship between protein levels and their corresponding mRNA levels in the DLPFC. We share our data in an integrative, searchable companion website to enable the discovery and localization of RNAs and proteins of interest for further investigation and to enhance our understanding of the temporally-defined molecular mechanisms governing typical and pathological DLPFC development.

## Materials and Methods

### Postmortem brain sample ascertainment

The current study analyzed fresh frozen postmortem dorsolateral prefrontal cortex (DLPFC) tissue (BA46) from 69 individuals varying in age from 39 days to 49.5 years (**Table S1**). The age range investigated in the current study reflects the vulnerability period for the development of neurodevelopmental and neuropsychiatric disorders. All samples were obtained from the National Child Health and Human Development Brain and Tissue Bank for Developmental Disorders at the University of Maryland, Baltimore, USA (UMBB). All subjects were defined as healthy individuals by forensic pathologists at UMBB, having no history of psychiatric or neurological complaints, also confirmed by next of kin interviews. These collected DLPFC samples comprised a broad range of developmental milestones, spanning neonatal (*n*=11), infantile (*n*=14), toddler (*n*=10), school aged (*n*=9), adolescence (*n*=8), young adulthood (*n*=9) and adulthood (*n*=8). Each developmental stage was matched for gender, postmortem interval (PMI), pH and ethnicity. The total sample included 41 males and 28 females covering African American (n=36), Caucasian (n=34), and Hispanic (n=1) ethnic backgrounds. The average measures for PMI and pH are as follows: PMI (17.1 ± 7.0 hrs.); pH (6.6 ± 0.2). A subgroup of these samples also underwent microarray transcriptome profiling (*n*=44), consisting of 27 males and 17 females, and covariates were recorded: pH (6.7 ± 0.15), PMI (16.9 ± 7.5 hrs.) and ethnicity (24 Caucasian, 20 African American). Detailed demographic and technical information on all samples can be found in **Table S1**.

### Tissue and protein extraction

Samples were dissected using a fine dental drill from the middle frontal gyrus at a level just rostral to the genu of the corpus callosum and the resulting tissue (average weight ~500 mg) was stored at ‐80°C until use. Proteins were extracted by sonicating each sample (~70mg) in 350ϋl lysis buffer (7M urea, 2M thiourea, 4% 3-[(3-cholamidopropyl)dimethylammonio]-1-propanesulfonate (CHAPS), 2% ASB14 and 70mM dithiotreitol (DTT), followed by sonication for 2 cycles of 15 seconds on ice using a Branson Sonifier 150 (Thistle Scientific; Glasgow, UK). A 50μl aliquot of each protein extract was precipitated for 4hr with 200ϋl of acetone at - 20°C. The precipitates were centrifuged for 30 min at 13400 rpm, at 4°C, the acetone supernatant decanted and discarded. The resulting pellets were re-suspended in 200ϋl of 50mM ammonium bicarbonate (pH 8.0) and sonicated for 10 seconds. Once re-suspended, protein concentration was measured by Bradford Assay. An aliquot of each precipitated protein extract, equivalent to 100ug of protein, was reduced with 100mM DTT for 30 min at 60°C, alkylated with 200mM iodoacetamide at room temperature for 30 min in dark and digested with 4ϋl of 0.5ϋl/ul of modified sequencing grade trypsin at 37°C for 17hrs. Digestion reaction was stopped by adding 0.80ϋl 8.8M Hydrochloric acid (1:60). An aliquot of 5μl from each digested sample was pooled together to be used as standard^26^.

### Liquid chromatography-mass spectrometry and protein quantification

Tryptic peptides were analyzed by a shotgun LC-MS approach using a 1290 Infinity LC coupled to Agilent 6550 iFunnel Q-TOF instrument (Agilent technology, USA). Peptide separation was carried out using an Agilent AdvanceBio Peptide column (2.1 μm x 250 mm, 2.7 ϋlm) over a 90 min linear gradient of 3 to 45% ACN. The flow rate was 0.3mL/min and the column temperature was set to 50°C. Peptides were then detected by quadrupole time-of-flight (Q-TOF) MS operated in positive mode. Acquisition was in data-dependent mode over m/z 300-1700. The top 10 precursor ions were scanned from 300-1700 and MS/MS from 50-1700. The precursor ions were then automatically isolated and fragmented using collision induced dissociation (CID) with a relative collision energy calculated using the formula, 3.6*(m/z)/100+- 4.8. Data files were processed by Spectrum Mill Protein Identification software (Rev B.05.00.180, Agilent Technologies, USA). The protein identification was executed against the Swiss-Prot database (released in February 2015, Homo sapiens). Search parameters were as follows; precursor mass tolerance, 20 ppm; product ion mass tolerance, 50 ppm; maximum two missed cleavages allowed; digested by trypsin; fixed modification of carbamidomethyl cysteine; variable modifications of oxidized methionine. After MS/MS searching, auto-validation was carried out by calculating the false-discovery rate (FDR). A FDR threshold of 1.2 was applied. Relative protein quantification was achieved using only distinct peptides that assigned to each protein. Unique peptide intensities were calculated from extracted ion chromatograms (MS1) from the precursor ions. Total peak intensities of all distinct peptides were then calculated to form relative protein expression levels.

### RNA isolation and microarray hybridization

All RNA procedures have previously been described^27^. Briefly, total RNA was extracted from dorsolateral prefrontal cortex samples using Trizol (Sigma-Aldrich, St. Louis, MO, USA) and RNA quality was assessed using a high-resolution electrophoresis system (Agilent Technologies, Santa Clara, CA, USA). Isolated total RNA was subjected to Affymetrix preparation protocol and each sample was hybridized to one HG-U133 Plus 2.0 GeneChip (Affymetrix, Santa Clara, CA, USA) to quantify transcriptome-wide gene expression.

### Data pre-processing

All data pre-processing and statistical analyses were conducted in the statistical package R. Proteins detected in at least 60% of all samples were labeled high-confidence proteins and used for downstream analyses. First, all data were normalized to fit approximate normal distribution. Protein data were median scaled by all runs and log(e) transformed. Protein Uniprot IDs were converted to HGNC symbols using the Uniprot database (http://www.uniprot.org/uploadlists/). Microarray data were normalized using the robust multiarray average normalization with additional GC-correction (GCRMA)^28^. When multiple microarray probes mapped to the same HGNC symbol, the probe the highest average expression across all samples was used. Following normalization, all data were inspected for outlying samples using unsupervised hierarchical clustering (based on Pearson coefficient and average distance metric) and principal component analysis to identify potential outliers outside two standard deviations from these grand averages; no outliers were present in these data. Linear mixed models from the R package^29^ were used to characterize and identify biological and/or technical drivers that may affect the observed RNA and protein abundance. This approach quantifies the main sources of variation in each expression dataset attributable to differences in age, age group, gender, PMI, pH and ethnicity. Finally, to identify age-related genes and proteins, generalized linear models with Bonferroni multiple test correction were implemented. The covariates gender, pH, PMI and ethnicity were included in the models to adjust for their potential confounding influence on RNA and protein expression (lm(Age ~ Expression + PMI + sex + pH + ethnicity)). Further, a Spearman’s correlation test was used to identify individual genes and proteins whose expression profile were significantly correlated with a pre-defined developmental stage or template (*e.g*. toddlers), which had been binarized Prior to network analysis, missing protein values were imputed using predictive mean matching in the MICE package^30^ (number of multiple imputations, *m*=5; the number of iterations, *maxit*=50). A high confidence set of proteins detected in at least 60% of the sample were used to make meaningful imputations. Weighted gene correlation network analysis (WGCNA) ^31^ was used to build signed co-expression networks independently for the transcriptome (*n*=20,122 genes) and proteome (*n*=584 proteins). To construct each network, the absolute values of Pearson correlation coefficients were calculated for all possible gene pairs (transcriptome data) and protein pairs (proteome data), and resulting values were transformed with an exponential weight (β) so that the final matrices followed an approximate scale-free topology (R^2^). Thus, for each network we only considered powers of β that lead to a network satisfying scale-free topology (*i.e*. R^2^>0.80), so the mean connectivity is high and the network contains enough information for module detection. The dynamic tree-cut algorithm was used to detect network modules with a minimum module size set to 30 and cut tree height set to 0.9999. The identified RNA and protein modules were inspected for association to age, as well as seven distinct postnatal stages and all recorded covariates. To do so, singular value decomposition of each modules expression matrix was performed and the resulting module eigengene (ME), equivalent to the first principal component, was used to represent the overall expression profiles for each module per sample. Modules were evaluated both quantitatively and qualitatively for expression patterns significantly associated with age (**Figure S3**). Fisher’s exact tests were used to assess the overlap of RNA and protein modules and correlations amongst RNA and protein ME’s were explored using Pearson’s correlation coefficients.

A series of module preservation analyses sought to determine whether (*i*) co-regulated modules of proteins are preserved at the RNA level and (*ii*) whether RNA modules are reproducible in independent BrainSpan data. We collected publically available BrainSpan data (http://www.brainspan.org/) and used only postnatal samples (*n*=17) to best reflect the developmental biology of our current sample (**Fig. S8**). For these analyses, module preservation was assessed using a permutation-based preservation statistic, *Z*_*Summary*_, implemented within WGCNA with 500 random permutations of the data^32^. *Z*_*summary*_ takes into account the overlap in module membership as well as the density and connectivity patterns of genes within modules. A *Z*_*summary*_ score <2 indicates no evidence of preservation, 2<*Z*_*summay*_<10 implies weak preservation and *Zsummary* >10 suggests strong preservation.

### Functional annotation and protein-protein interaction networks

All age-related RNAs and proteins identified through either linear regression or network-based analyses, were subjected to functional annotation using the ToppFun module of ToppGene Suite software^33^. We explored gene ontology terms related to biological processes and molecular factors using a one-tailed hyper-geometric tested (Benjamini-Hochberg FDR corrected) to assess the significance of the overlap. All terms must pass an FDR corrected p-value and a minimum of five genes/proteins per ontology were used as filters prior to pruning ontologies to less redundant terms.

The STRING database v9.1^34^ was used to assess whether RNA and protein modules were significantly enriched for direct protein-protein interactions (PPIs) and to identify key genes/proteins mediating the regulation of multiple targets. For these analyses, our signature query of RNA or protein modules were used as input. STRING implements a scoring scheme to report the confidence level for each direct PPI (low confidence: <0.4; medium: 0.4-0.7; high: >0.7). We used a combined STRING score >0.04. Hub genes within the PPI network are defined as those with the highest degree of network connections. We further used STRING to test whether the number of observed PPIs were significantly more than expected by chance using a nontrivial random background model. For visualization, the STRING network was imported into Cytoscape^35^.

### Cell type and genetic risk loci enrichment analyses

CNS cell type specific markers were collected from three independent resources, including cell type specific genes from RNA-sequencing^36,37^ and mass spectrometry-based proteomics^38^. In order for a gene/protein to be labeled cell type specific, each marker required a minimum log2 expression of 1.4 units and a difference of 0.8 units above the next most abundance cell type measurement, as previously shown^18^. Mouse homologues were identified and converted into human HGNC gene symbols using the mygene R package^39^. In parallel, neurodevelopmental disorder genetic risk loci were curated from human whole exome and genome-wide association studies of autism spectrum disorder^40^, epilepsy^41^, developmental delay (OMIM)^42^, intellectual disability^15^ and schizophrenia^43^. Overrepresentation of cell type markers and genetic risk-related gene sets within proteome and transcriptome modules was analyzed using a one-sided Fisher exact test to assess the statistical significance. All P-values, from all gene sets and modules, were adjusted for multiple testing using the Benjamini Hochberg procedure. We required an adjusted P-value <0.05 to claim that a gene set is enriched within a module. Lists of neurodevelopmental genetic risk loci can be found in **Table S7**.

### Cell type deconvolution

The frequencies of brain cell types were estimated for proteomic using Cibersort^44^ cell type deconvolution (https://cibersort.stanford.edu/). Cibersort relies on known cell subset specific marker genes and applies linear support vector regression, a machine learning approach highly robust compared to other methods with respect to noise, unknown mixture content and closely related cell types. As input, we used a curated cell type specific protein signature matrix^38^ to distinguish between neurons, oligodendrocytes, astrocytes and microglia. We were unable to obtain sufficient enough overlap for microglial markers based on protein detection to make meaningful predictions for this cell type.

### Data availability

To promote the exchange of this information, we developed an interactive website with an easily searchable interface to act as a companion site for this paper: the DEveLopmental Trajectory Atlas (DELTA) in DLPFC is available from the following URL: http://amp.pharm.mssm.edu/DELTA. In addition, all sample descriptions, proteomic and transcriptomic data are available and can directly downloaded from this site. Alternatively, gene expression data can be downloaded from GEO using accession GSE13564.

## Results

### Proteome organization in the developing dorsolateral prefrontral cortex

A shotgun proteomics approach was applied to measure temporal protein abundance in 69 DLPFC samples (Brodmann area 46) from normal individuals, aged 39 days to 49.5 years of age. Following standardized data pre-processing (**Fig. S1**), we detected 911 proteins for which a total of 386 proteins were assigned low-confidence measures of protein detection and 584 proteins were assigned high-confidence on the basis of being detected across >60% of all samples (see methods). For the low-confidence proteins, rates of protein detection were moderately influenced by different developmental stages, whereby 60 proteins, which were predominately post-synaptic density proteins, were more likely to be detected during early developmental stages and 99 proteins, which were implicated in cellular respiration and GTPase binding, were more likely to be detected during adulthood (**Fig. S2**, **Table S2**).

To reduce the probability of false positives, we restricted downstream analyses to only proteins with high-confidence levels of protein detection. For these proteins, a substantial amount of protein expression variation was explained by age relative to other biological factors (**Fig. 1 AC**). Next, we sought to identify proteins that were significantly regulated as a function of postnatal age and identified 83 proteins (FDR *p*<0.05), including 66 with decreasing abundance and 17 with increasing abundance across postnatal stages (**Table S2**). The top ten most significantly increased and decreased age-related proteins are displayed in **Table 1**. Several significant age-related proteins mapped to known neurodevelopmental genetic risk loci, including genetic loci implicated in autism spectrum disorder (ASD) (*ANK2*, *YWHAE*, *L1CAM*, *FABP5*, *NRCAM*), intellectual disability (*L1CAM*, *PLP1*, *PSAP*, *QDPR*) and schizophrenia (*NCAN*, *ALCAM*, *GNAO1*, *PSAP*, *NFASC*). In addition, we also observed that the neonatal time period (38-89 days) explained the largest fraction of protein level variability according to age, including 131 neonatal-related proteins (FDR p<0.05) strongly enriched for central nervous system development, neurogenesis and gliogenesis (**Fig. S3**, **Table S3**). Notably, no proteins displayed sex-dependent effects across development.

**Figure 1.**
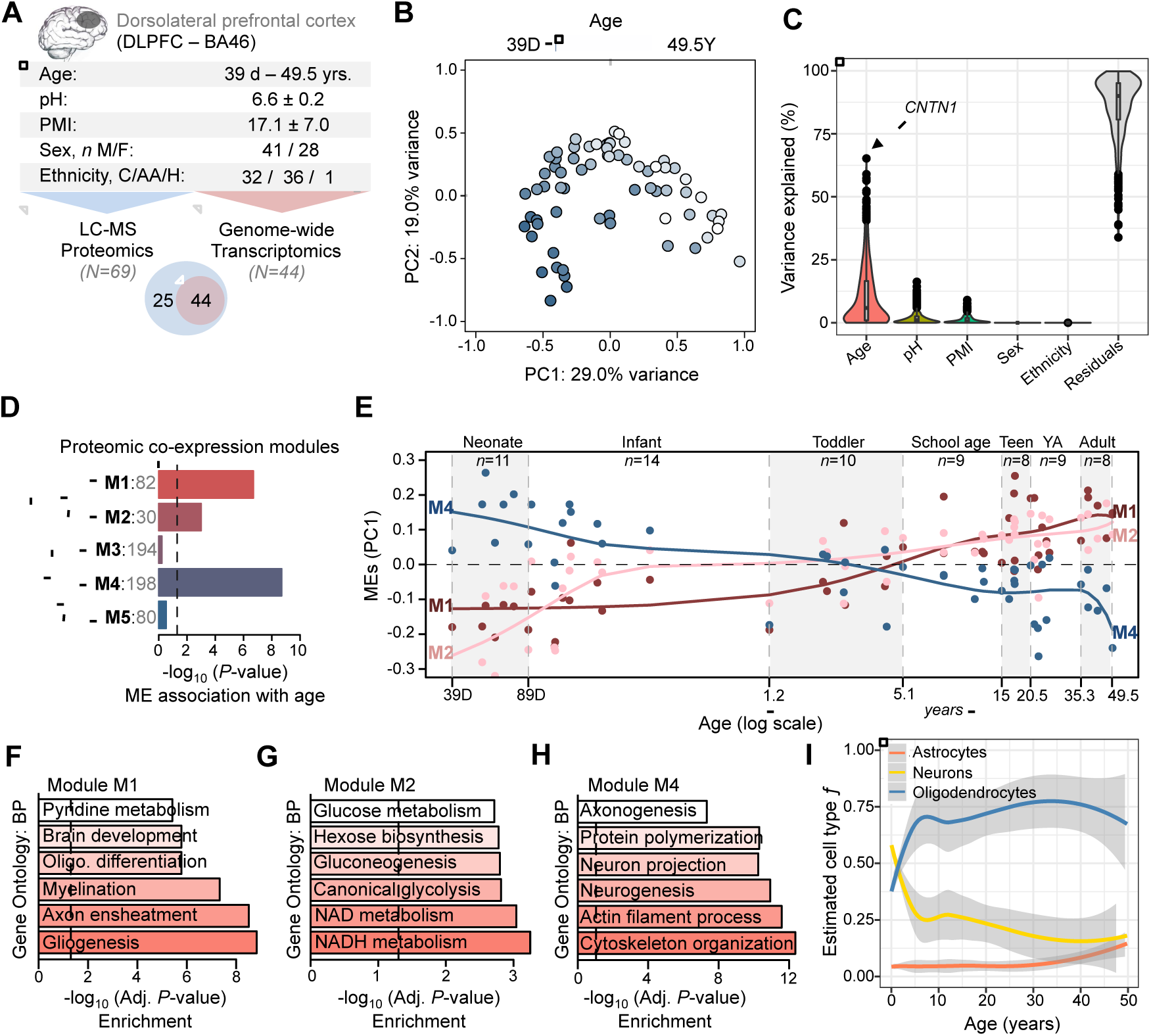
Protein expression and function in the developing DLPFC. (**A**) Sample characteristics for LC-MS proteomic data (*N*=69) and a subgroup of DLPFC samples with additional transcriptome data (*N*=44). (**B**) Principal component analysis on global normalized protein abundance, samples are shaded by age (39 days - 49.5 years). (**C**) variancePartition linear mixed model analysis of global protein abundance identifies age as a leading trait explaining most of observed protein variability. (**D**) Weighted correlation network analysis identified five protein modules, three significantly associated with age. X-axis indicates ‐log_10_ P-value significance. Module names and the number of proteins within each module are displayed right of dendrogram. (**E**) Age (x-axis) relative to module eigengene (ME) expression (y-axis) for modules M1 and M2 increasing throughout development and module M4 decreasing throughout development. Smoothing linear spline models (knot=4) were fit for each ME across age. Seven post-natal developmental stages are color shaded and labeled (SA, school age; YA, young adult). Functional annotation of age-related modules M1 (**F**), M2 (**G**) and M4 (**H**) according to gene ontology biological processes. (**I**) Cell-type deconvolution on global protein abundance. **Data information:** In (**C**) HGNC symbol abbreviations; *CNTN1*, contactin 1; *HNRNPU*, heterogeneous Nuclear ribonucleoprotein U; *CSRP1*, cysteine and glycine rich protein 1; *CLTB*, clathrin light chain b; *COX7C*, cytochrome c oxidase subunit 7c.

**Table 1.** Top ten up and down age-related proteins throughout postnatal development with paired RNA products.

| Uniprot ID | HUGO gene symbol (name) | Proteome |  | Transcriptome |  |
| --- | --- | --- | --- | --- | --- |
|  |  | *t*-statistic | Adj. *P*-value | *t*-statistic | Adj. *P*-value |
| O15075 | *DCLK1* (doublecortin like kinase 1) | -10.786 | 6.02E-16 | -3.836 | 4.46E-04 |
| P22676 | *CALB2* (calbindin 2) | -9.154 | 3.51E-13 | -11.731 | 2.31E-14 |
| Q9BPU6 | *DPYSL5* (dihydropyrimidinase like 5) | -8.951 | 7.86E-13 | -12.522 | 3.05E-15 |
| P52306 | *RAP1GDS1* (Rap1 GTPase-GDP dissociation stimulator 1) | -8.950 | 7.90E-13 | -5.331 | 4.39E-06 |
| P05413 | *FABP3* (fatty acid binding protein 3) | -8.511 | 4.57E-12 | -2.295 | 2.72E-02 |
| P29966 | *MARCKS* (myristoylated alanine rich protein kinase C substrate) | -8.321 | 9.79E-12 | -11.097 | 1.24E-13 |
| Q12860 | *CNTN1* (contactin 1) | -7.733 | 1.05E-10 | -3.490 | 1.22E-03 |
| P55072 | *VCP* (valosin containing protein) * | -7.688 | 1.25E-10 | -3.995 | 2.78E-04 |
| Q6PCE3 | *PGM2L1* (phosphoglucomutase 2 like 1) | -7.639 | 1.53E-10 | -7.150 | 1.33E-08 |
| O14594 | *NCAN* (neurocan) ‡ | -7.256 | 7.13E-10 | -8.759 | 9.49E-11 |
| P09417 | *QDPR* (quinoid dihydropteridine reductase) † | 7.707 | 1.16E-10 | 10.303 | 1.09E-12 |
| P49189 | *ALDH9A1* (aldehyde dehydrogenase 9 family member A1) | 6.201 | 4.81E-08 | 2.478 | 1.77E-02 |
| O94856 | *NFASC* (neurofascin) ‡ | 5.917 | 1.47E-07 | 3.699 | 6.65E-04 |
| P07602 | *PSAP* (prosaposin) † ‡ | 5.818 | 2.16E-07 | 9.804 | 4.47E-12 |
| P12277 | *CKB* (creatine kinase B) | 5.590 | 5.22E-07 | N/A | N/A |
| O94811 | *TPPP* (tubulin polymerization promoting protein) | 5.257 | 1.86E-06 | 10.842 | 2.47E-13 |
| P14618 | *PKM* (pyruvate kinase, muscle) | 5.190 | 2.39E-06 | 10.344 | 9.73E-13 |
| Q16653 | *MOG* (myelin oligodendrocyte glycoprotein) | 5.180 | 2.48E-06 | 4.991 | 1.29E-05 |
| P13611 | *VCAN* (versican) | 5.170 | 2.58E-06 | -2.601 | 1.31E-02 |
| Q969P0 | *IGSF8* (immunoglobulin superfamily member 8) | 5.038 | 4.21E-06 | 2.104 | 4.19E-02 |

Linear regression models adjusted for pH, PMI, sex and ethnicity. The direction of change (*t*-statistic) across postnatal development and *P*-value significance are displayed for each protein and paired RNA product. Candidate neurodevelopmental genetic risk loci are symbolized: \*, autism; †, intellectual disability; ‡, schizophrenia. N/A indicates not detected.

To characterize the temporal organization of the proteome we ran unsupervised weighted correlation network analysis and identified five modules of co-regulated proteins, three of which were significantly associated with development and age (**Fig. 1D**, **Fig. S4-5**). Two age-related modules displayed increasing expression throughout development (FDR p<0.05) and were enriched for processes related to myelination and gliogenesis (M1, 82 proteins, *r*=0.71, *p*=1.0×10^-11^) and gluconeogenesis and NADH metabolism (M2, 30 proteins, *r*=0.58, *p*=2.0×10^-7^) (**Fig. 1 E-G**). One age-related module displayed decreasing expression across development and was implicated in axonogenesis, neurogenesis and cytoskeleton organization (M4, 198 proteins, *r*=-0.77, *p*=2.3×10^-14^) (**Fig. 1H**). The remaining two modules were not significantly associated with age nor with any other developmental stages or technical factors, and were enriched for functions related to cellular respiration and ATP metabolic processes (M3, 194 proteins) and nucleotide metabolism and oxidation-reduction processes (M5, 80 proteins) (**Fig. S6**). Collectively, all five modules were significantly enriched for direct protein-protein interactions (PPI), beyond what would be expected by chance (**Table S4**), and PPI networks were constructed for each module (**Fig. S6-7**). Densely connected hub proteins for each age-related module included, *MBP* and *PGM1* for module M1, *ENO2* and *MDH1* for module M2 and *UBC* and *HSP90AA1* for module M4. Further, cell type deconvolution revealed that the majority of proteins expressed across all DLPFC samples were specific to neuronal and oligodendrocyte cell types, and further highlighted substantial decreases in neuronal cell populations paralleled by increases in oligodendrocytes throughout development (**Fig. 1I**); results which correlated with transcriptome-based estimates (**Fig. S8**).

### Correspondence between transcriptome and proteome module organization

A total of 556 common HGNC symbols were detected between our high confidence, reproducible proteins and transcriptome-wide gene expression assays (**Fig. 2A**). Similar to the proteome, the largest amount of gene expression variability was explained by age, as compared to any other factor (**Fig. S1**). Comparably, the neonatal time period (39-89 days) also explained the largest fraction of transcriptome variation by age, albeit to a lesser extent than in the proteome, including genes primarily involved in ATP metabolic processes (**Fig. S2**). Meanwhile, variation across the sexes was small genome-wide, but it explained a large percentage of expression variation for genes on chrX and chrY. Linear regression analyses identified ~11.5% of the transcriptome was associated with postnatal development, including 1145 genes with increasing expression and 1181 genes with decreasing expression across all developmental stages (FDR *p*<0.05). (**Fig. S2**, **Table S5**). These age-related genes also included several known neurodevelopmental genetic risk loci implicated in ASD (*LRP1*, *RNF135*, *YWHAE)*, schizophrenia (*TEKT4*, *LRP1*, *DNAH1*, *BRSK1*, *INTS1*, *ZC3H10*, *METTL14*) and developmental delay (*SCYL1*, *PIGQ*, *OBSL1*, *SMARCB1*, *CEP135*, *SPG11*, *TAF1*, *TAT*, *FAM126A*, *RAD21)*. Notably, 27 molecules were uniquely detected at the protein level, and a significant fraction were enriched for oxidative phosphorylation-related processes (FDR *p*=2.3×10^-8^).

**Figure 2.**
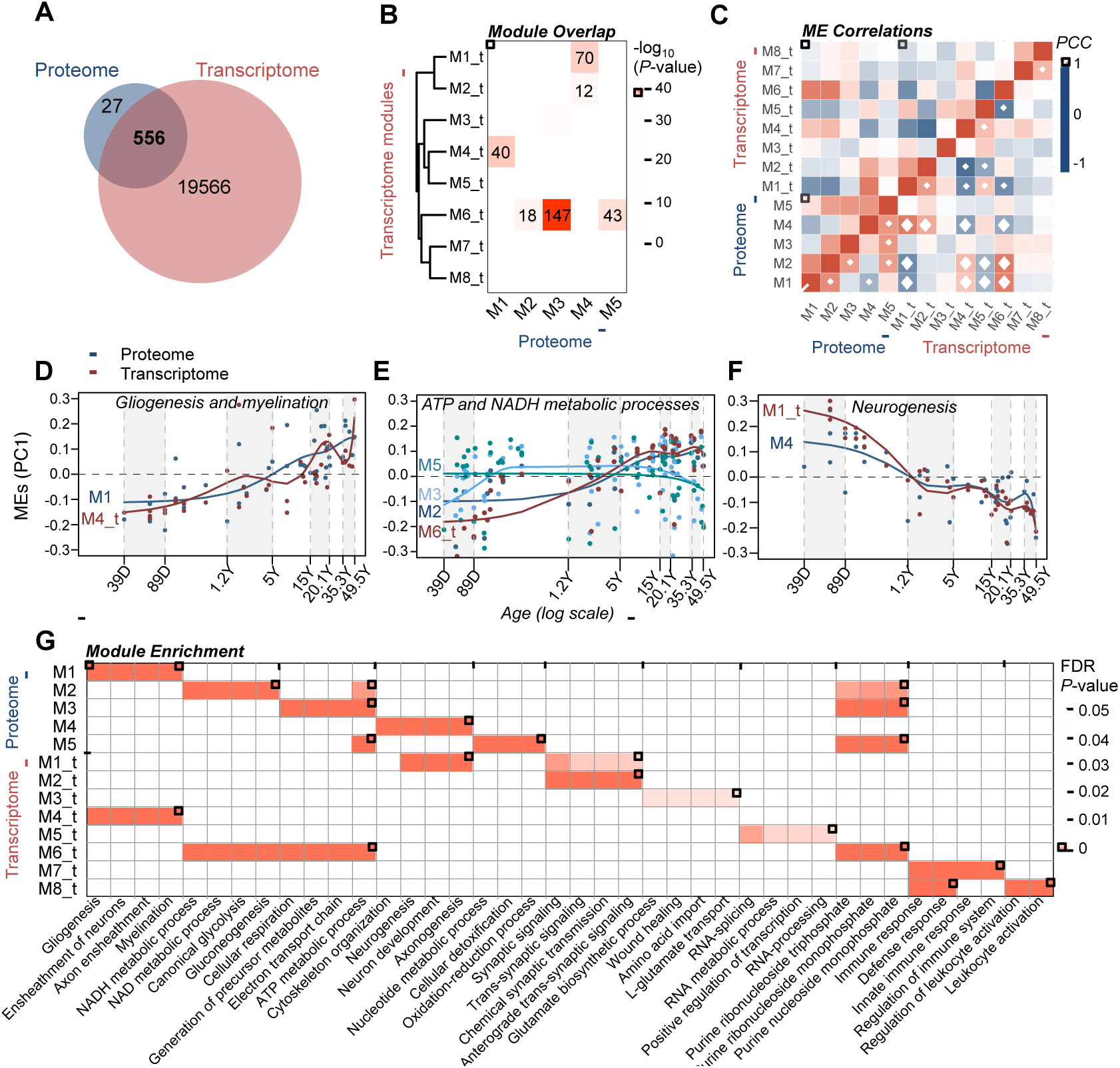
Overlap of age-related RNA and protein modules. (**A**) Venn diagram of detected the proteome and transcriptome. (**B**) Overlap analysis of transcriptome‐ and proteome-based modules. The number of overlapping HGNC symbols are displayed for significant intersects. (C) Pearson’s correlation coefficient analysis of module eigengenes (MEs) within and across transcriptome‐ and proteome-based modules. White diamonds indicate *p*<0.05. Convergent ME expression patterns for modules with significant overlap and shared function between transcriptome and proteome, including gliogenesis and myelination (**D**), ATP and NADH metabolic processes (**E**) and neurogenesis (**F**). Smoothing linear spline models (knot=4) were fit for each ME across seven postnatal stages (shaded grey/white). (**G**) Overlap and correspondence of the top four functional categories for all RNA‐ and protein-based modules.

Weighted correlation network analysis identified eight modules of co-regulated genes (**Fig. 2B**), which displayed a high degree of reproducibility compared to existing postmortem BrainSpan data (**Fig. S9**). Four of the eight modules were significantly associated with postnatal development, including two modules implicated in gliogenesis (M4_t, 2624 genes, *r*=0.54, *p*=2.4×10^-4^) and ATP metabolic processes (M6_t, 5078 genes, *r*=0.72, *p*=4.1×10^-5^) with increasing expression, and two modules involved in neurogenesis (M1_t, 3046 genes, *r*=-0.71, *p*=8.6×10^-8^) and synaptic signaling (M2_t, 403 genes, *r*=-0.53, *p*=2.2×10^-4^) with decreasing expression throughout postnatal development (**Fig. S10**). One immune response-related module was significantly associated to the toddler postnatal age group (1.2-5.1 years) (M7_t, 384 genes, *r*=0.52, *p*=3.3×10^-4^). Interestingly, a number of significant overlaps were identified between transcriptome modules and proteome modules (**Fig. 2B**). The majority of overlapping RNA-and protein-based modules also displayed a high degree of collinearity (**Fig. 2C**) and shared similar biological functions, including modules involved in gliogenesis and myelination (M1, M4_t), ATP metabolic processes (M2, M3, M5, M6_t) and neurogenesis (M4, M1_t) (**Fig. 2D-G**). Overall, all five protein modules were well represented at the RNA level (**Fig. 2G**).

### Correspondence between RNA and protein levels throughout postnatal development

We further quantified the association between gene and protein level expression using a subgroup of DLPFC samples for which paired transcriptome and proteome data were available (*n*=44). We first examined the degree of within-sample correlation using 556 paired RNAs and proteins, and then by sub-setting these analytes according to the five previously identified protein modules (**Fig. 3A**). Across all analytes, we found weak-to-moderate within-sample correspondence (Pearson’s *r*=0.15-0.40), with the highest correlations observed for RNA and protein products involved in gliogenesis and myelination (M1, *r*=0.26-0.53) and the lowest correlations for RNA and protein products involved in nucleotide and ATP metabolic processes (M5, *r*=-0.05-0.22). Subsequently, we explored these within-sample correlations as a function of age and found that the correspondence between RNAs and their respective translated proteins is higher for early developmental stages and lower for later developmental stages, indicating an overall decrease in correlation between RNA and protein level expression throughout postnatal development (*r*=-0.56, *p*=6.4×10^-5^). This decreased correlation was also significant for subgroupings of RNAs and proteins involved in processes of myelination and gliogenesis (M1, *r*=-0.30, *p*=0.04) and neurogenesis and cytoskeleton organization (M4, *r*=‐ 0.67, *p*=3.9×10^-7^) (**Fig. 3B,E**). These age-related differences prompted us to investigate the degree of conservation in protein-based co-expression networks at the RNA level, for which we found no preservation for co-regulated RNAs implicated in nucleotide metabolic processes (M5) (**Fig S11**). Finally, we examined correlations between the 556 RNA-protein pairs across all samples (opposed to within-samples), as a function of postnatal development. Overall, a high level of correspondence was observed when comparing age-related linear regression results (¿-statistics) computed separately for individual RNAs and proteins (*r*=0.62, *p*=2.4×10^-60^) (**Fig. 3C**). Upon closer inspection, 78.4% of all RNA-protein pairs were positively correlated throughout development while the remaining were negatively correlated (**Table S6**). Notably, we find that RNA expression profiles that are significantly associated with cortical development (FDR *p* < 0.05) display higher correlations with their respective translated protein level expression compared to RNA expression profiles that are not significantly age-related (*p*=6.4×10^-25^) (**Fig. 3D**).

**Figure 3.**
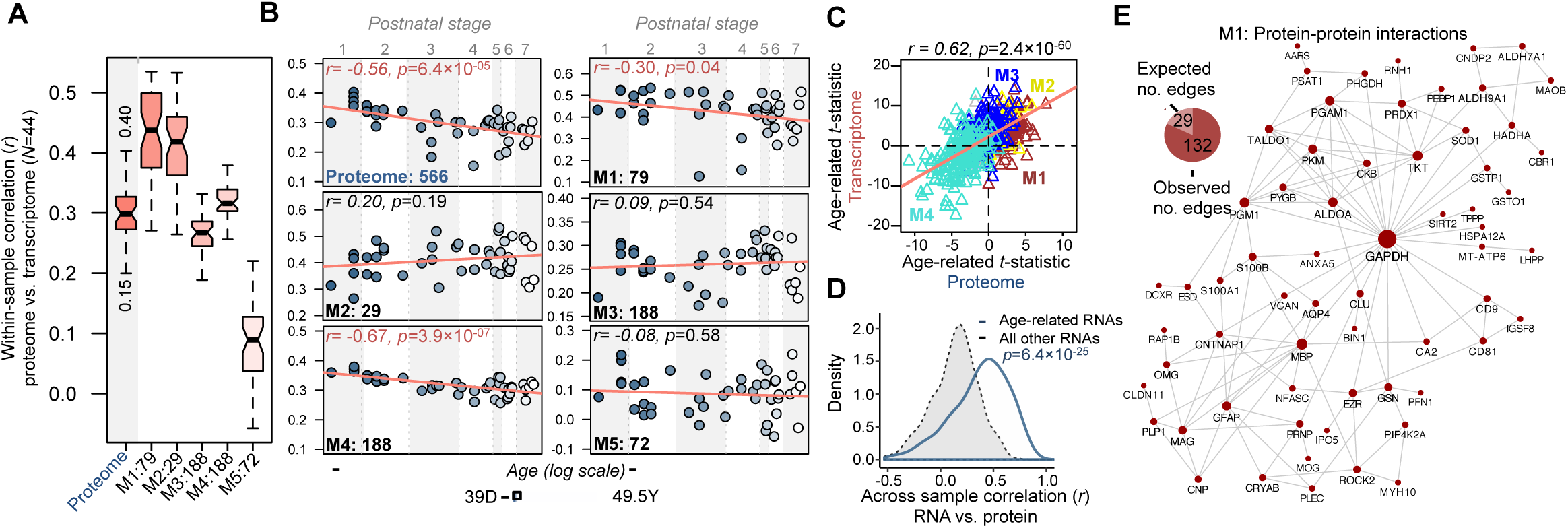
Concordance between transcriptome and proteome in the developing DLPFC. (**A**) *Within-sample* Pearson correlation coefficients (PCC) between paired transcriptome and proteome samples (*N*=44). Associations were tested for all paired RNA and protein molecules (proteome) and then by focusing on RNA and protein content accordingly to protein module status. The number of molecules compared within each module are listed on the x-axis. (**B**) Average PCC’s within paired transcriptome and proteome samples (y-axis) measured as a function of age (x-axis). Samples are ranked according to age, and shaded accordingly to postnatal developmental period (1, neonate; 2, infant; 3, toddler; 4, school age; 5, teenager; 6, young adult; 7, adult). (**C**) Scatterplot of age-related linear regression t-statistics computed for the overlapping 556 mRNAs and proteins, colored according to protein module membership. (**D**) *Across-sample* Spearman correlation coefficient (SCC) comparing RNA expression profiles to their respective translated protein expression. SCC’s are parsed by RNA’s which are significantly associated with cortical development (FDR p <0.05), compared to those which do not (**Table S6**). A Mann-Whitney U test was used to compute significance between these two groups. (**E**) Direct protein-protein interaction network of protein module M1, displaying significant decrease in correlation between RNA and proteins across postnatal development.

### Cell type and neurodevelopmental disorder genetic risk loci enrichment

We sought to determine whether the transcriptome‐ and proteome-based modules were strongly linked to the underlying cellular architecture in the developing DLPFC using previously defined cell type specific markers (**Fig. 4A**). Three different cell type specific resources were used to discover and validate cell type enrichments, including those based on RNA^36,37^ and protein discovery^38^. As expected, several proteome and transcriptome modules were significantly enriched for known cell type specific markers and demonstrated high reproducibility across three independent resources. Protein module M1 was consistently enriched for oligodendrocyte and astrocyte cell types. Protein modules M3 and M4 also consistently displayed significant over-representation for neuronal cell type markers. In parallel, several transcriptome-based modules displayed consistent enrichment for CNS cell type markers, including modules M1_t, M2_t and M6_t enriched for neuronal markers, M3_t enriched for astrocyte markers, M5_t enriched for oligodendocyte markers, and M7_t enriched for microglial markers. Notably, no protein module displayed enrichment for microglial cell markers, which is consistent with our cell type estimates that indicate our DLPFC proteome samples are predominately comprised of neuronal and oligodendocytes cell types.

**Figure 4.**
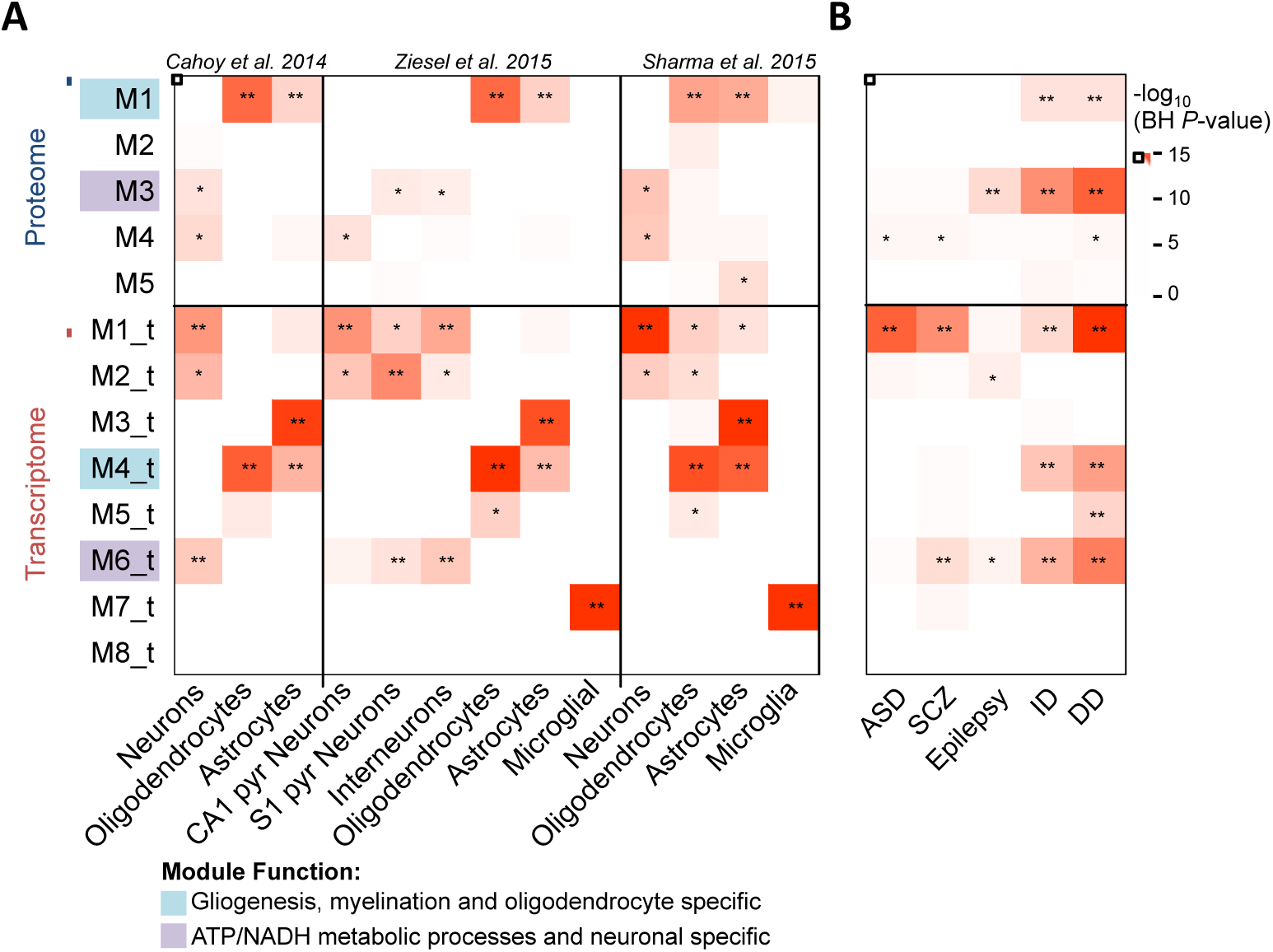
Cell type and neurodevelopmental disorder genetic risk loci enrichment. (**A**) Cell type enrichment analysis for the identified transcriptome‐ and proteome-based modules. The number of significantly overlapping HGNC symbols are displayed (**, *p*<0.0001; *, 0.01 > *P* < 0.05). (**B**) Genetic risk loci enrichment analysis according to transcriptome‐ and proteome-based modules. Five previously generated lists of neurodevelopmental disorder risk loci were used, including intellectual disability (ID), developmental delay (DD), autism spectrum disorder (ASD), epilepsy and schizophrenia (SCZ). **Data information:** In (**A**) Three cell type marker resources were leveraged, including two based on transcriptome profiling^36,37^and one based on proteome profiling^38^.

Subsequently, we sought to determine whether genes associated with risk for neurodevelopmental disorders converge on common cellular and biological processes during human cortical development in the proteome and transcriptome. Intellectual disability (ID) genes tightly coalesce in proteome and transcriptome modules that implicate gliogenesis (M1, ⋂=7, *p*=0.009; M4_t, ⋂=70, *p*=0.006) and ATP metabolism functions (M3, ⋂=20, *p*=2.2×10^-9^; M6_t, ⋂=120, *p*=0.01) (**Fig. 4B**). Similarly, developmental delay (DD) risk variants were concentrated in the same modules associated with gliogenesis (M1, ⋂=11, *p*=0.001; M4_t, ⋂=147, *p*=0.003) and ATP metabolism (M3, ⋂=35, *p*=2.4×10^-12^). The ID genes used here represent high confidence genes implicated in monogenic forms of ID from multiple publications, whereas the DD genes are those available from the Developmental Disorders Genotype-Phenotype Database (DDG2P). Both ID and DD are relatively common pediatric disorders with overlapping symptomologies. These loci also constituted several hub proteins within proteome module M3, including *NDUFS1-8*, *SYN1*, *STXBP1* and *MAP2K1* (**Fig S6A**).Together, these results support the notion that common and rare variants contribute to ID and DD by perturbation of processes encoding ATP metabolism and myelination. In addition, similar to previous reports, we also confirm strong enrichment for several ASD and SCZ genetic risk loci in neurogenesis-related module M4 in the proteome (⋂=4, *p*=0.04; ⋂=10, *p*=0.04, respectively) as well as module M1_t in the transcriptome (⋂=59, *p*=8.2×10^-16^; ⋂=134, *p*=4.3×10^-9^, respectively). These analyses, and others, can be performed using our online software tool (http://amp.pharm.mssm.edu/DELTA).

## Discussion

Proteins are the functional components of cells in the CNS, however our understanding of the brain proteome continues to lag behind the pace of transcriptome discovery. This discrepancy is largely due to the lack of established proteome-wide technologies, which have only recently matured to enable improved protein detection and coverage. To provide a foundation for an age-dependent brain proteome map, we performed label-free LC-MS proteomic analysis across 69 human DLPFC (BA46) samples, aged 35 days to 49.5 years of age, which comprised seven different developmental stages. Our approach identified 911 highly abundant and reproducible proteins across a large number of developmentally distinct biological replicates and resulted in the largest collection thus far of protein expression data in the developing human DLPFC. The proteins detected here function to sub-serve some of the most fundamental CNS cell signaling cascades required for typical cortical development. By integrating these data with transcriptome data, we were able to examine relationships between RNA and protein expression levels, which revealed a much tighter coupling of RNA and protein expression during early developmental stages (*i.e*. neonatal and infant) compared to later stages (*i.e*. adulthood). Finally, we examined RNA and protein networks enriched for neurodevelopmental genetic risk loci, to gain insight into how mutations in risk genes may perturb molecular pathways during healthy brain development. We discuss these points in turn below.

The majority of the detected proteins in the current investigation (~64.1%) were associated with cortical development and formed functional protein modules, which harbored a large number of direct protein-protein interactions. Two protein modules were identified, which gradually increased in expression across cortical development and enriched for gliogenesis, myelination and olidodendrocyte cell type specificity (M1) as well as NADH metabolism and gluconeogenesis (M2), while one module was decreasing in expression and implicated in axonogenesis, cytoskeleton organization and neuronal cell types. These modules represent some of the most basic CNS functions and their expression profiles are at least partially driven by a shifting CNS cellular landscape throughout cortical development, as reflected by the observed neuron-glia oscillations. Although cell division and migration of neurons are largely prenatal events, neurogenesis is known to persist throughout adult life, albeit to a limited level and produce only a small fraction of the neuronal population^46,47^. In contrast, proliferation and migration of glial progenitors, while beginning prenatally, continue for a protracted period as oligodendrocytes and astrocytes differentiate. Oligodendrocyte cells begin to differentiate by increasing myelin protein expression, as evident in the current study. However, much uncertainty has existed regarding the extent of postnatal proliferation, migration and differentiation, and about the timing of these processes relative to each other^46^. Our results indicate that the greatest degree of change likely occur during school age years (3-15 years of age), and that these neuron-glia changes appear to play an important role in the functional organization of neural circuits during early and late stages of postnatal development. We also report marked increases in discretely co-regulated proteins involved in NADH metabolism and gluconeogenesis (M2) across development, which is consistent with the well-known energy requirements of the brain^48^. Previous work by us and others suggests that myelination is also a major energy-demanding process in the brain^27,49^, especially during postnatal life. For example, myelin synthesis is an ATP-dependent process and oligodendrocytes often oxidize glucose at higher rates than neurons^50^, supporting these distinct changes in protein modules across time. To this end, two additional protein modules were identified peaking in expression during the ages of 6 months to 1 year and were enriched for cellular respiration and ATP metabolism (M3) and purine ribonucleoside monophosphate activity (M5), which likely represent shared components of a larger glycolysis, cellular respiration and oxidative phosphorylation cycle, along with M2. Importantly, glucoregulatory abnormalities, oxidative stress vulnerability and oligodendrocyte dysfunction have been prominently linked to neuropsychiatric and neurodegenerative disorders^51-53^, and a detailed understanding of how these proteins unfold in expression throughout cortical development may guide future follow-up studies targeting these pathways.

In the context of the temporally dynamic expression profiles, a fundamental question is whether RNAs and their respective translated proteins correlate throughout postnatal development. We observed within-sample Pearson correlation coefficients between 0.15 and 0.40. Several studies have also found similar low correlations in human^19,20^ and murine tissues^38^. These discrepancies may be due to well-known differences in the regulation, localization, structures and functions of mRNA and proteins. However, a novel finding from the current study, is that when presenting these correlations as a function of postnatal age, we identified that correlations between RNA and protein expression tend to decrease throughout development (*r*=-0.56, *p*=6.4*10), in that younger samples tend to have higher RNA-protein correlations and older samples tend to have weaker RNA-protein correlations (**Fig 3B**). This negative trend accelerated for genes implicated in myelination (M1) and cytoskeleton organization (M4). Interestingly, the efficiency of myelination decreases with age, a process largely regulated by age-dependent epigenetic control of gene expression^54^. That is, during infancy, myelin synthesis is preceded by down-regulation of oligodendrocyte differentiation inhibitors, and this is associated with recruitment of histone deacetylase to promoter regions; a process that becomes less efficient in adulthood and ultimately prevents a successive surge in myelin gene expression^54-56^. Regarding the weakening correlation of cytoskeleton-related RNA and protein expression across postnatal development, it is clear that cytoskeleton plays a vital role in regulating CNS cell mechanics with age. Moreover, since several studies have shown that many neurodevelopmental disorders are likely influenced by aberrant cytoskeleton organization, it is important to understand how the expression and interaction of cytoskeletal proteins change with age. Overall, these results shed light on several candidate myelination and cytoskeleton proteins for follow-up functional studies to assess whether insufficient amounts of translated protein product during early development may negatively impact nerve cell shape, motility and communication thereby leading to behavioral and/or developmental deficits.

We also examined the correspondence between 556 RNA-protein pairs across all samples and found that the majority of pairs (78.4%) correlate positively across development, while others do not, and in some instances display strong negative correlations (**Table S6**). It is unlikely that false positives can fully explain these low/negative correlations. Therefore, it may be that varied levels of regulation, such as translational regulation, supersede the transcriptional level and provide biological fine-tuning for the specific conditions encountered by the cells. Furthermore, protein half-life and translational rates can also vary, which can effect the correlation between RNA and protein levels. However, it is notable that RNA profiles, which were not associated with postnatal development displayed significantly lower correlation distributions compared to RNA profiles, which were significantly associated with aging and development (**Fig. S11**). Thus, our results show that significant age-related changes in gene expression commonly co-occur with tighter correlations with protein levels, giving further confidence for the use of mRNA data for biological discovery.

One important similarity across the brain transcriptome and proteome was the consistent mapping of intellectual disability and developmental delay genetic risk loci to modules enriched for myelination, gliogenesis, and ATP metabolic processes. These modules displayed a collinear patter of expression between RNA and protein products, peaking in expression during adolescence and adulthood (**Fig. 2D-F**). As there is a close interdepency between myelin synthesis and ATP-dependent processes, a disorder affecting one of the two inevitably also leads to disturbance of the other. Indeed, defective myelination and ATP processes have been reported as key factors causing pathogenic processes involved in these disorders^15,57,58^, and these data provide further substantial evidence in the broader context of long-term brain development. These results support the notion that common and rare variants contribute to ID and DD by perturbation of common gliogenesis and ATP metabolism networks. In parallel, ASD variants resided primarily in neuronal-based modules in the transcriptome, and not in the proteome. These results echo recent large transcriptome network studies that demonstrate the involvement of neuronal and synaptic processes involved in ASD^12,16,17^. We also mapped epilepsy genetic risk loci to oxidative phosphorylation-related modules in the proteome, consistent with growing evidence that deficits in oxidative phosphorylation complexes can result in increased oxygen and free-radical release likely implicated in the initiation and progression of epilepsy^59^.

Our study also has some limitations. First, while our selective approach sought to inform CNS development by detecting highly abundant and reproducible proteins across 69 developmentally distinct biological replicates, the data presented here may represent an incomplete picture of the entire proteome. We are unable to discuss the developmental role of lowly abundant proteins. Moreover, 556 proteins were represented at the mRNA level, a marginal 3.5% of the detected transcriptome. Despite, with this level of detection we were able to capture a considerable amount of protein variability and functionality across postnatal development compared to paired transcriptome data. That is, all of the age-related transcriptome modules (M1_t, M2_t, M4_t, M6_t: 11,150 genes total), which comprised 55.4% of the observed transcriptome, were well represented at a functional level in the proteome, even though fewer proteins were detected; emphasizing that the majority of the detected proteins are highly expressed and sub-serve for some of the most fundamental CNS molecular processes. Nonetheless, it is possible that future reports applying deeper analytical techniques will enable both greater proteome coverage. A second caveat to these data is the lack of prenatal samples, developmental stages when gene expression patterns appear to be most dynamic. For example, vast increases in expression for synapse and dendrite development genes occur prenatally and taper off in the first decade of postnatal life^10,60^, and went undetected in the current investigation likely due to the lack of prenatal samples. Moreover, it is challenging to directly compare these age-related proteomics results to those derived from other studies due to extensive differences in proteomic technologies and the ascertainment of postmortem tissues. However, in contrast to previous proteomic studies, we were able to capitalize on larger postnatal developmental group sizes (2.5x larger), thus increasing our ability to identify biologically meaningful age-related proteins and protein networks. As these concerns are addressed in the future, it will be possible to reveal further insights into the transcriptional and translational foundations of human brain development.

Our unbiased, global approach outlined both similarities and differences of the developing DLPFC between the transcriptome and proteome across postnatal development. The various proteins detected and discussed are likely to be candidates for further functional and/or synaptic developmental studies. Therefore, to promote the exchange of this information, we developed a website with an easily searchable interface to act as a companion to this resource paper: the DEveLopmental Trajectory Atlas (DELTA) in DLPFC is available from the following URL: http://amp.pharm.mssm.edu/DELTA. This website will be maintained and periodically updated as additional data emerge from this unique cohort. Using the website researchers can: *1)* query any protein/gene symbol of interest to determine at which developmental stage it is expressed; *2)* determine whether a user submitted input list of proteins/genes is overrepresented within our identified proteomic and transcriptomic gene modules; *3)* download all corresponding proteomic and transcriptomic data. Our expectation is that the website and the data that it hosts will serve as a resource to stimulate and enable additional studies to further elucidate the complex molecular controls guiding postnatal human cortical development.

## Acknowledgements

This study was supported by the Stanley Medical Research Institute. M.S.B. is funded by the Autism Science Foundation (#17-001) and the Seaver Autism Center for Research and Treatment at the Icahn School of Medicine at Mount Sinai. A.M. and Z.W. are partially supported by NIH grants U24CA224260 and U54HL127624. M.G.G is supported by a Gonville & Caius College/Cambridge Home and European Scholarship Scheme (CHESS) EU Maintenance Bursary and an EPSRC Doctoral Training Grant (DTG) studentship. C.S.W is a recipient of a National Health and Medical Research Council (Australia) Principal Research Fellowship (PRF) (#1117079).

## Conflict of Interest

S.B. is a director of Psynova Neurotech Ltd and PsyOmics Ltd. C.S.W is on an advisory board for Lundbeck, Australia Pty Ltd and in collaboration with Astellas Pharma Inc., Japan.

## Author Contributions

S.O. and S.B. conceived the study and designed the experiments. M.J.W. and C.S.W. selected and obtained the brain tissue samples. M.G.G. and N.R. prepared the samples for proteomics analysis. Z.W., M.S.B. and A.M. developed the DELTA in DLPFC website. S.O. and J.M.R. pre-processed the data. M.S.B. processed and analyzed the data. M.S.B. and S.O. wrote the manuscript with input from S.B. and J.D.B. M.J.W. and C.S.W. also provided critical input regarding data interpretation and editing of manuscript. All other co-authors read and approved the final manuscript.

